# In-Cell Protein Crystallization via a Locally Flexible 24-mer Assembly Precursor

**DOI:** 10.64898/2026.08.25.746898

**Authors:** Satoshi Abe, Junko Tanaka, Kosuke Kikuchi, Tadaomi Furuta, Yuta Aizawa, Yoshikazu Tanaka, Takeshi Yokoyama, Shuji Kanamaru, Ririko Kobayashi, Takafumi Ueno

## Abstract

In-cell protein crystallization (ICPC) produces ordered protein crystals within living cells, but the mechanisms used by proteins to acquire long-range crystalline order in the cellular environment remains poorly understood. Here, we define the assembly pathway of CipB, a crystalline inclusion protein from *Photorhabdus luminescens*. CipB crystals formed in cells dissolve under mild acidic conditions into a predominant 24-mer species, supporting a model in which an in-cell crystal is built from a discrete 24-mer assembly precursor rather than through direct packing of smaller oligomeric states. Structural analysis of recrystallized CipB shows that the same 24-mer architecture packs into a body-centered cubic lattice, consistent with the lattice observed for the in-cell crystals. Cryo-EM and molecular dynamics analyses indicate that the 24-mer assembly precursor preserves its overall architecture while retaining local conformational flexibility at the N-terminal and surface-loop regions. Mutation analyses further link the N-terminal region to the formation of the 24-mer precursor and surface residues to lattice assembly. These observations support a stepwise crystallization model in which N-terminal flexibility facilitates the formation of an assembly-competent 24-mer precursor, whereas defined hydrophobic surface contacts subsequently organize these precursors into a long-range-ordered lattice.

## Introduction

In-cell protein crystallization (ICPC) occurs in diverse cellular compartments, including the cytoplasm, nucleus, endoplasmic reticulum (ER), and secretory granules.(1–7) Unlike conventional *in vitro* crystallization, which requires high-purity protein preparation and extensive screening of crystallization conditions, ICPC generates long-range crystalline order spontaneously within the crowded and heterogeneous environment of living cells.(1–4) Recently, in-cell protein crystals have been explored as platforms for structural biology, including in-cell crystallographic pipelines and *in situ* electron diffraction analysis.(8–12) Molecular crowding, localized protein concentration, and liquid-liquid phase separation have been implicated in ICPC, yet the structural pathway through which soluble protein assemblies transition into long-range crystalline order remains poorly understood.(2, 6)

A well-characterized example of an in-cell protein crystal is the polyhedra crystal (PhC).(13, 14) PhC is a highly ordered cubic lattice crystal that is formed in insect cells infected with cytoplasmic polyhedrosis virus (CPV) to encapsulate viral particles.(13, 15) The crystal structure of PhC reported in 2007 revealed that the trimer of the polyhedra monomer is a building block, forming stable crystals through dense packing.(13) Insulin is another representative in-cell protein crystal which assembles at high concentrations within secretory granules and forms a crystal in a process promoted by binding of Zn^2+^.(16, 17) These examples have shaped a view of in-cell protein crystallization as the packing of defined, symmetric, and relatively stable protein assemblies. This view is further reinforced by recent advances in protein crystal design, in which rigid, pre-organized components with precisely engineered interfaces are used to construct ordered lattices.(18) However, it remains unclear if ICPC can also proceed through flexible assembly precursors, and how such precursors retain flexibility while acquiring long-range crystalline order.

CipB provides a useful system for addressing this question. CipB forms in-cell crystals in the entomopathogenic bacterium *Photorhabdus luminescens* and can also be crystallized in *E. coli*.(19, 20) Here, we define the crystallization pathway of CipB by integrating structural, computational, and mutational analyses. We show that CipB forms a soluble 24-mer assembly precursor that serves as the building block of the crystalline lattice. Flexible N-terminal interactions are positioned at inter-tetramer interfaces and support formation of the 24-mer precursor, whereas defined surface contacts mediate precursor packing into the ordered lattice. These findings support a model in which CipB crystallization proceeds through a locally flexible 24-mer assembly precursor rather than through direct crystallization of monomers, or rigid pre-organized assemblies.

## Results

### CipB crystals dissolve into a discrete 24-mer assembly in solution

To understand the molecular mechanism of CipB crystallization, we first examined the pH-dependent dissolution of CipB crystals and the oligomeric state of the resulting soluble species. In-cell protein crystals of CipB (**CipB_icc**) were produced in *E. coli*, as previously reported,(19) appearing as well-defined cubic crystals with an average size of 0.6 ± 0.1 μm (Figs. 1a and 1b). Matrix-assisted laser desorption/ionization time-of-flight mass spectrometry (MALDI-TOF MS) analysis of **CipB_icc** showed a peak of 11,319 Da, consistent with the calculated molecular weight of the CipB monomer (11,315.6 Da) (Fig. S1).

**Fig. 1.**
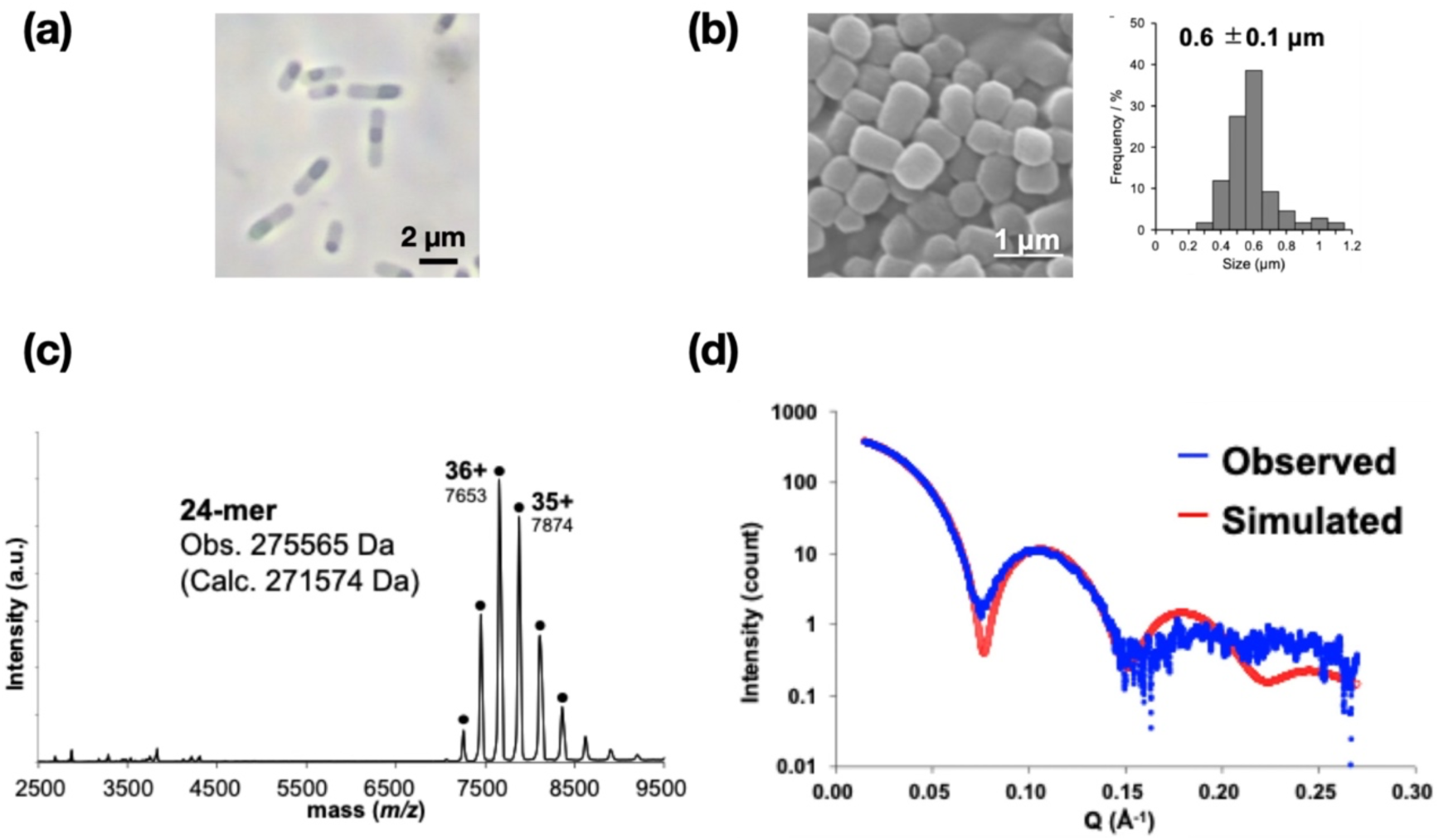
(a) Phase-contrast micrograph of *E. coli* expressing CipB, showing intracellular crystal formation. (b) SEM image and size distribution of purified **CipB_icc**. (c) ESI-TOF mass spectrum of **CipB_sol** in 100 mM ammonium acetate. (d) SAXS profile of **CipB_sol** in 100 mM NaOAc buffer at pH 5.0 (blue curve) fitted with a core-shell model using SasView (red curve).

To investigate the pH-dependent solubility of **CipB_icc**, crystals were immersed in buffers across a wide pH range (3–10). **CipB_icc** was found to exhibit high stability at pH 6.0-9.0, but solubility was found to increase under more acidic (pH 3.0-5.0) or basic (pH 10.0) conditions (Fig. S2a). Native-PAGE analysis of supernatant fractions showed two dominant bands at approximately 270 and 90 kDa under acidic (pH 3.0-5.0) and basic (pH 10.0) conditions (Fig. S2b). The ratio of high- to low-molecular-weight species varies with pH, with the larger assemblies predominating at pH 5.0, the mildest condition at which the crystals dissolve (78:22 ratio). Dynamic light scattering (DLS) measurements indicated an average hydrodynamic diameter of 10.5 ± 0.2 nm at pH 5.0. In comparison, smaller molecular species (5.5-6.0 nm) were observed at pH 3.0 and 10.0, suggesting partial dissociation into smaller oligomers (Fig. S2c). To characterize the main soluble species after dissolving the crystals at pH 5.0, the supernatant was purified by size-exclusion chromatography (SEC), which showed a single peak, indicating the presence of a major homogeneous assembly in solution (Fig. S3a). Electrospray ionization time-of-flight (ESI-TOF) mass spectrometry of the purified CipB (**CipB_sol**) revealed a major assembly corresponding to a 24-mer (observed: 275,565 Da, calculated: 271,574 Da) (Fig. 1c). Small-angle X-ray scattering (SAXS) analysis of **CipB_sol** revealed a scattering pattern characteristic of a hollow, near-spherical particle with outer and inner diameters of 10.4 and 5.4 nm, respectively (Fig. 1d), consistent with the hydrodynamic size measured by DLS. Native-PAGE analysis of **CipB_sol** showed a predominant band at approximately 270 kDa, consistent with the 24-mer assembly identified by ESI-TOF MS, with only a faint additional band at approximately 90 kDa (Fig. S3b). These results show that CipB predominantly exists as a discrete 24-mer cage-like assembly in solution at pH 5.0, suggesting that this assembly represents a plausible soluble precursor or repeating unit of the CipB crystal.

### CipB adopts a 24-mer assembly in a body-centered cubic crystal lattice

To define the structural relationship between the soluble 24-mer assembly and the crystalline lattice of CipB, we determined the crystal structure of CipB. A SAXS analysis of **CipB_icc** showed Bragg diffraction peaks consistent with a body-centered cubic (bcc) lattice with a lattice constant of 111.3 Å (Fig. S4). However, synchrotron diffraction analysis of **CipB_icc** at SPring-8 BL32XU did not yield diffraction data suitable for structure determination, likely because of the limited diffraction quality of the in-cell crystals. To overcome this issue, **CipB_sol** was recrystallized *in vitro*, yielding larger crystals (∼400 μm, Fig. 2a). The structure of recrystallized **CipB_sol** was determined at 3.5 Å resolution (Table S1). The crystals have a bcc lattice (*I*432 space group, *a* = *b* = *c* = 110.9 Å), which closely matches the lattice constant estimated for **CipB_icc** by SAXS. This indicates that CipB forms the same crystalline lattice architecture both *in vitro* and in cells (Fig. S4 and Table S1). The structure was determined by molecular replacement using the CipA structure (PDB: 7XHS) as a search model.(21) Each CipB monomer adopts a compact oligonucleotide/oligosaccharide-binding (OB) fold composed of five β strands (β1–β5) and one α-helix (α1), with β1–β3 and α1 located in the N-terminal half and β4–β5 in the C-terminal region (Fig. 2b). Despite having only 23% sequence identity, a structural comparison with CipA revealed high similarity (Cα RMSD of 1.8 Å), suggesting that CipB preserves a similar overall fold.(21) No electron density was observed for the N-terminal region (Met1–Ile13) and several loop regions (Leu32–Pro34, Leu53, and Lys86), suggesting conformational disorder or flexibility in these regions (Fig. 2b).

**Fig. 2.**
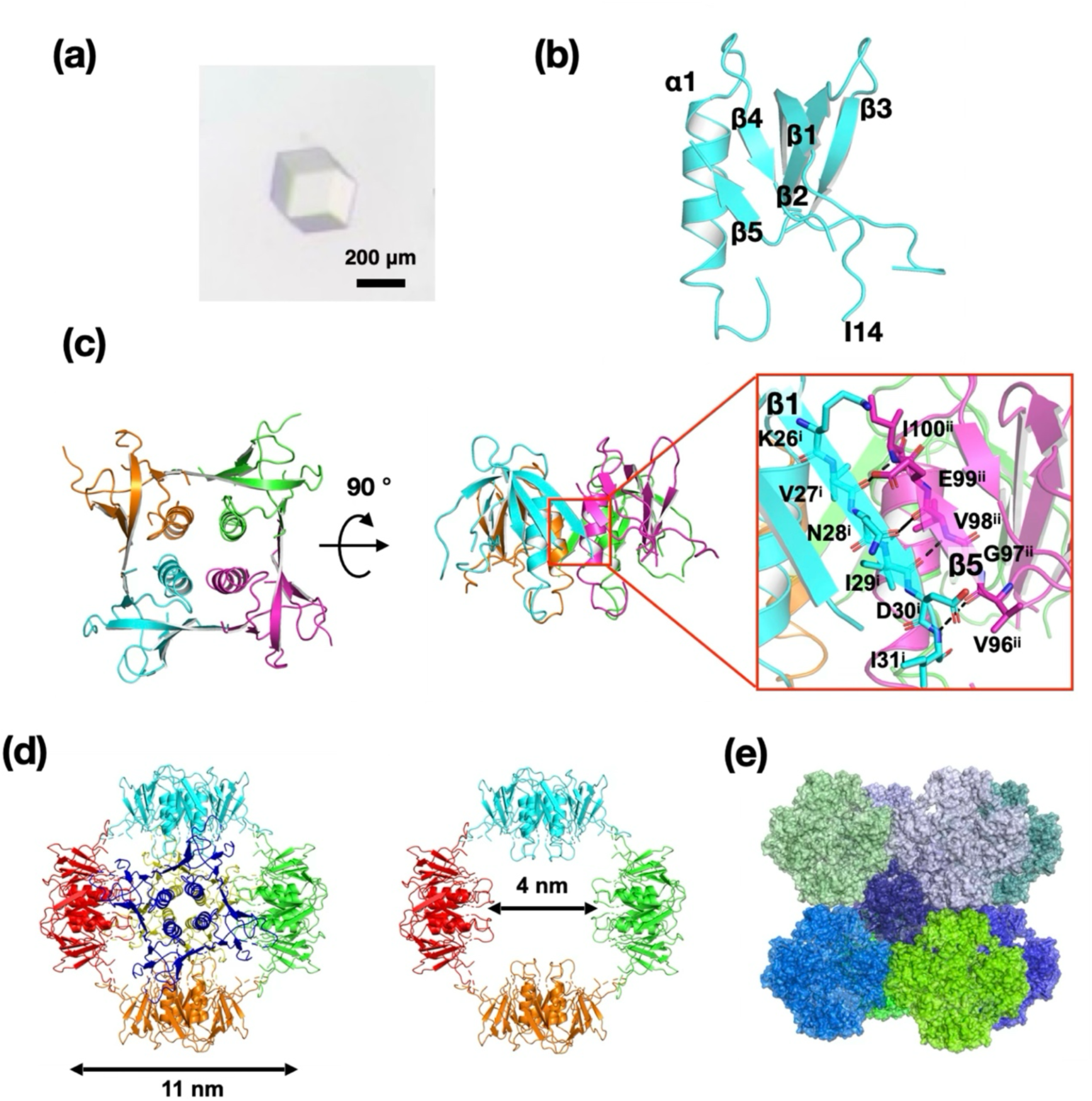
Crystal structure of recrystallized CipB. (a) An optical image of a CipB single crystal obtained by recrystallization. (b) Crystal structure of the CipB monomer. (c) Tetramer of CipB mediated by intermolecular β-sheet interactions. i and ii indicate different monomers within the tetramer. (d) Assembly of six tetramers into a 24-mer cage-like structure. (e) Crystal packing of CipB 24-mers forming a bcc lattice with space group *I*432.

CipB monomers assemble into tetramers through β-sheet formation mediated by hydrogen bonds (Fig. 2c). At the inter-monomer interface in the tetramer, β1 (residues 26-31) and β5 (residues 96-100) align to form an extended β-sheet, stabilized by a network of backbone-backbone hydrogen bonds (O/Val27^i^-N/Ile100^ii^, N/Ile29^i^-O/Val98^ii^, O/Ile29^i^-N/Val98^ii^, and N/Ile31^i^-O/Val96^ii^) and backbone-sidechain contacts (O_δ_/Asn28^i^-O/Val98^ii^ and O/Val27^i^-O_ε_/Glu99^ii^).

In the crystal lattice, these tetramers are symmetrically arranged to form a 24-mer cage-like assembly, with outer and inner diameters of 11 and 4 nm, respectively (Fig. 2d). The 24-mer cages are further organized into a highly ordered body-centered cubic (bcc) lattice in space group *I*432. This suggests that the 24-mer assembly represents the repeating structural unit of **CipB_icc** (Fig. 2e). Within this 24-mer assembly, strong continuous inter-tetramer contacts were not clearly resolved, consistent with the observation of conformational heterogeneity in the flexible regions, including the N-terminus. The first interface between 24-mers within the lattice involves side-chain contacts between adjacent cages, such as Val46^a^-Val46^b^, Leu44^a^-Val46^b^, and Val46^a^-Leu44^b^ (Fig. 3a). The second interface involves the His75–Leu78 region (Fig. 3b). These localized cage-cage contacts may help explain the observation of the periodic arrangement of CipB 24-mer assemblies in the bcc lattice and the preservation of the overall architecture of each assembly.

**Fig. 3.**
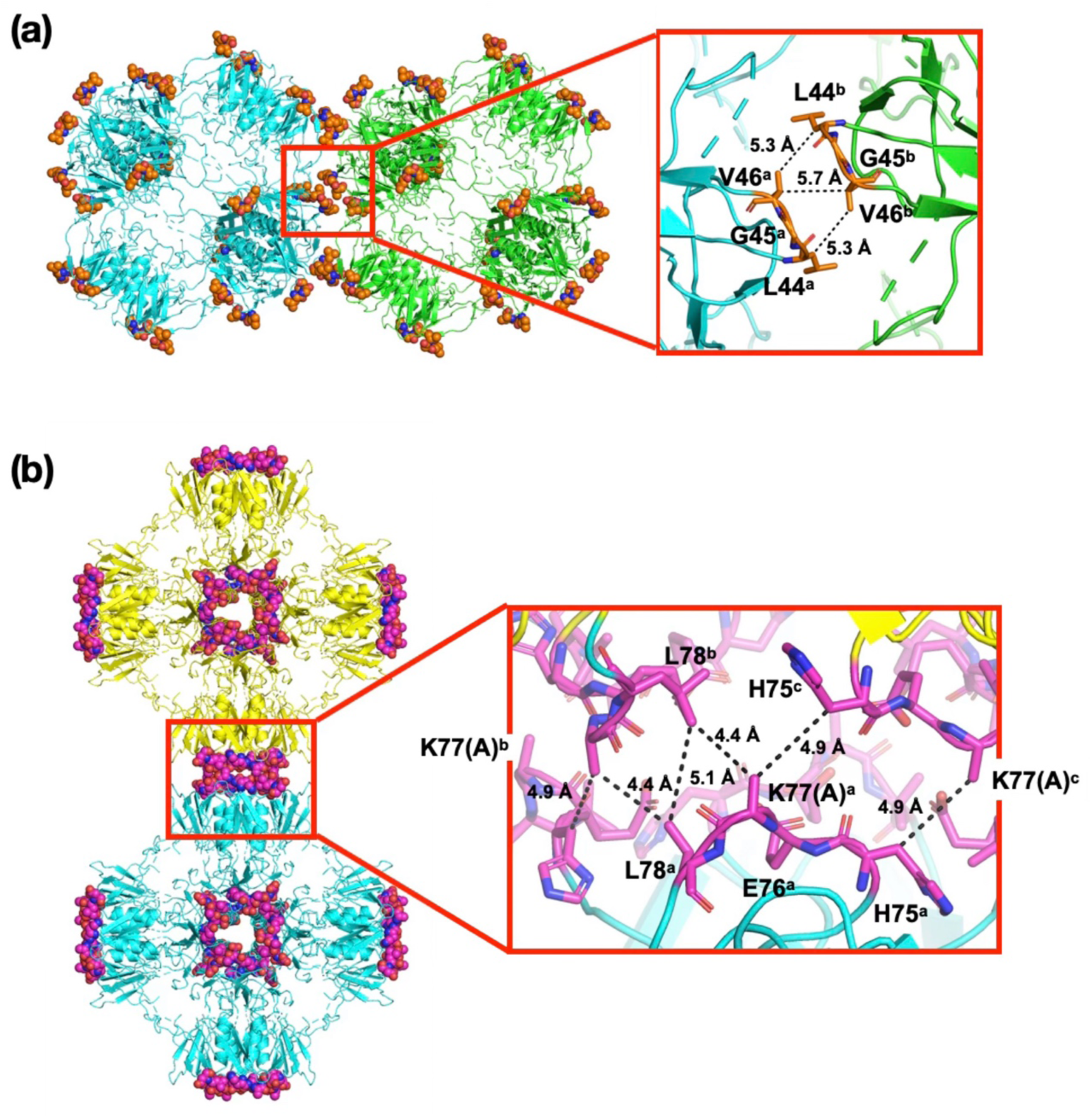
Intermolecular interactions mediating CipB crystal lattice formation. (a) Hydrophobic contacts involving the Leu44-Val46 loop, and (b) interface stabilized by interactions between residues His75-Leu78. Due to the lack of electron density, the side chain of Lys77 was modeled as Ala in the structural model. Residues from different 24- mer cages are labeled as a, b, and c to indicate their respective origins within the lattice.

### Cryo-EM structure of the CipB 24-mer assembly in solution

To investigate the solution-state structure of the CipB 24-mer assembly, **CipB_icc** was dissolved in 100 mM sodium acetate buffer at pH 5.0 and the resulting supernatant was subjected to single-particle cryo-electron microscopy (cryo-EM) analysis. A three-dimensional reconstruction map was obtained at a resolution of 3.6 Å from approximately 1.7 million particles (Figs. 4a, and S5). The atomic model of CipB, determined by X-ray crystallography, was fitted and refined into the cryo-EM map, revealing a solution-state structure closely resembling the crystallographic 24-mer assembly (Fig. 4b).

**Fig. 4.**
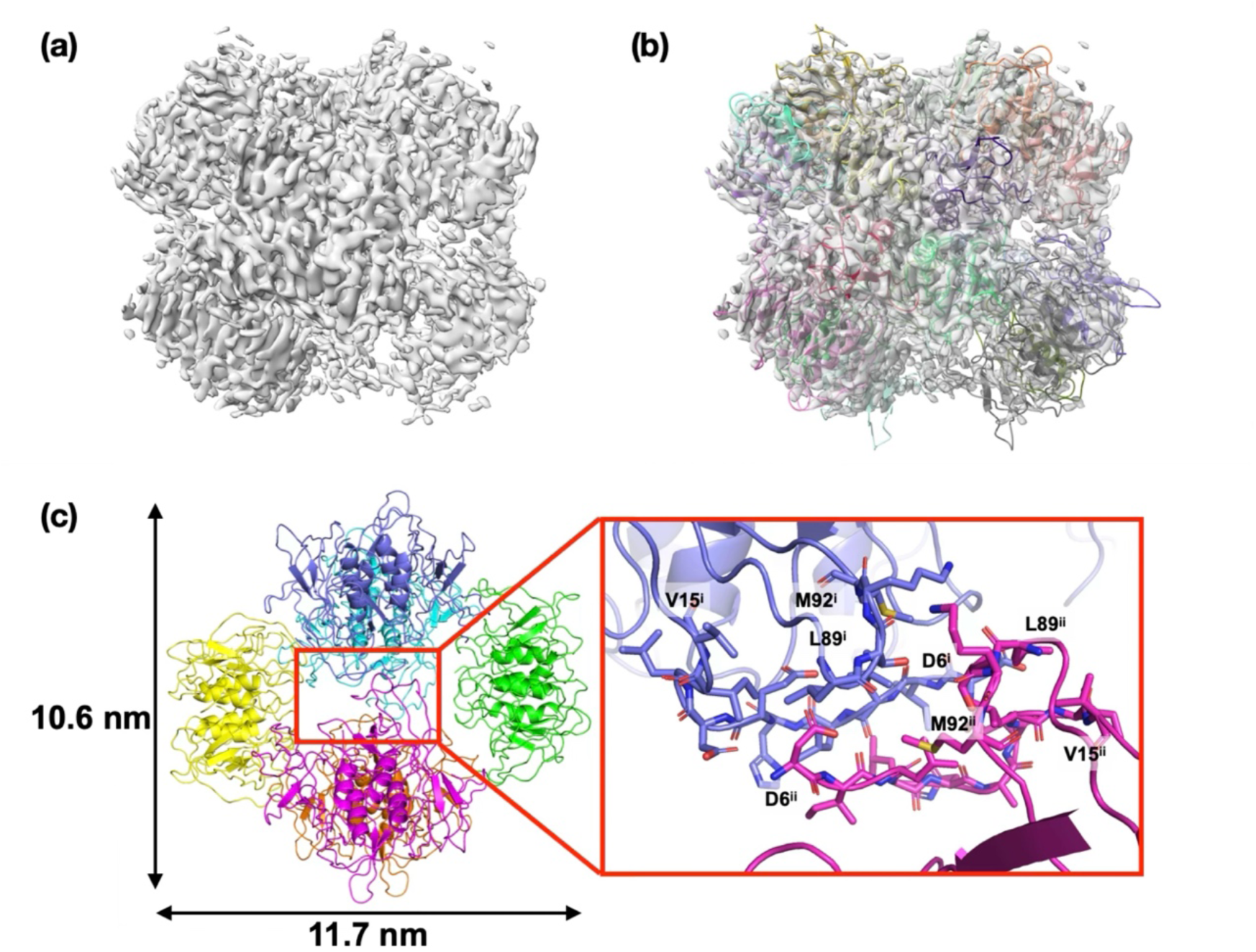
Cryo-EM structure of the CipB 24-mer cage. (a) The cryo-EM reconstruction map of CipB 24-mer cage at 3.6 Å resolution. The cage exhibits a near-spherical geometry with internal voids consistent with the crystallographic structure. (b) Atomic model of the CipB 24-mer fitted into the cryo-EM map. (c) Enlarged view of the interface between adjacent tetramers, mediated by the N-terminal region (Asp6-Val15) and the loop region (Leu89-Met92), suggesting the involvement of these regions in mediating inter-tetramer interactions within the 24-mer cage.

The cryo-EM analysis provided structural information on flexible regions which are disordered in the crystal structure. Cryo-EM maps corresponding to the N-terminal region (Asp6–Val15), which was not resolved in the crystal structure, were observed at seven of the twelve tetrameric interfaces within the 24-mer assembly (Fig. 4c). At these interfaces, the N-terminal regions are positioned to face each other across adjacent tetramers and are located near the Leu89–Met92 region (Fig. 4c). In addition, map density corresponding to surface-exposed loop regions, including residues Leu44-Val46 and His75-Lys77, was found to be weak or poorly resolved in many monomers within the 24-mer assembly. To further compare the solution and crystalline states, we calculated per-residue Cα RMSDs between the cryo-EM and crystal structures (Fig. S6). The analysis revealed structural deviations in the N-terminal region and several loop regions located on the cage surface (Leu44-Val46), within the cage interior (Ser35 and Leu54-Asp56), and at the inter-tetramer interface (Asn88-Leu89), whereas the β-sheet-rich core was found to remain highly conserved.

The overall 24-mer assembly was found to exhibit an anisotropic cage-like morphology relative to the crystallographic model (Fig. 4). The diameters measured across two orthogonal cross-sections are 11.7 and 10.6 nm, respectively, corresponding to an approximately 10% difference and indicating a clear deviation from spherical symmetry (Fig. 4c). Furthermore, map density in some of the tetrameric units was found to be weaker than in others, suggesting uneven local order within the 24-mer assembly.

In addition to the 24-mer assembly (∼10 nm), 2D classification identified a minor population of smaller particles (∼5 nm) (Figs. S3 and S5f). However, higher-resolution reconstruction of these particles was not possible because of their small sizes and structural heterogeneity. These observations indicate that the CipB 24-mer assembly is retained in solution but is not completely uniform at the local structural level.

### MD simulations of 24-mer (cage) and 8-mer (dimer-of-tetramers) CipB

To explore the stability and structural dynamics of the 24-mer cage of CipB, we performed three independent 500-ns molecular dynamics (MD) simulations (Fig. 5). In all three runs, the overall cage structure was maintained, although the relative orientations of the six tetramers were found to fluctuate slightly (Fig. 5a, Movie S1). The root mean square deviations (RMSDs) from the initial structure are approximately 6 Å (Fig. 5b), and the distances between opposing tetramers remained at approximately 80 Å throughout the simulations (Fig. 5c). Analysis of the root mean square fluctuations (RMSFs) revealed that RMSF values at the N-terminus of each monomer exceeded ∼7 Å, indicating significant flexibility in these regions (Fig. 5d). To examine possible interactions between the N-terminal region and the disordered loop regions observed in the X-ray crystal structure, we analyzed contacts involving Leu53 in the final MD structures at 500 ns (Fig. 5e). Leu53 was found to frequently contact residues in the Lys4-Ile14 region, particularly Leu8 and Leu13 (Fig. 5f). Other disordered regions, including Leu32-Pro34 and Lys86, were also found to contact N-terminal residues during the simulations (Fig. S7a). These observations suggest that the flexible and hydrophobic N-terminal region interacts with multiple loop regions and may contribute to the local structural disorder observed for Leu32-Pro34, Leu53, and Lys86 in the X-ray crystal structure.

**Fig. 5.**
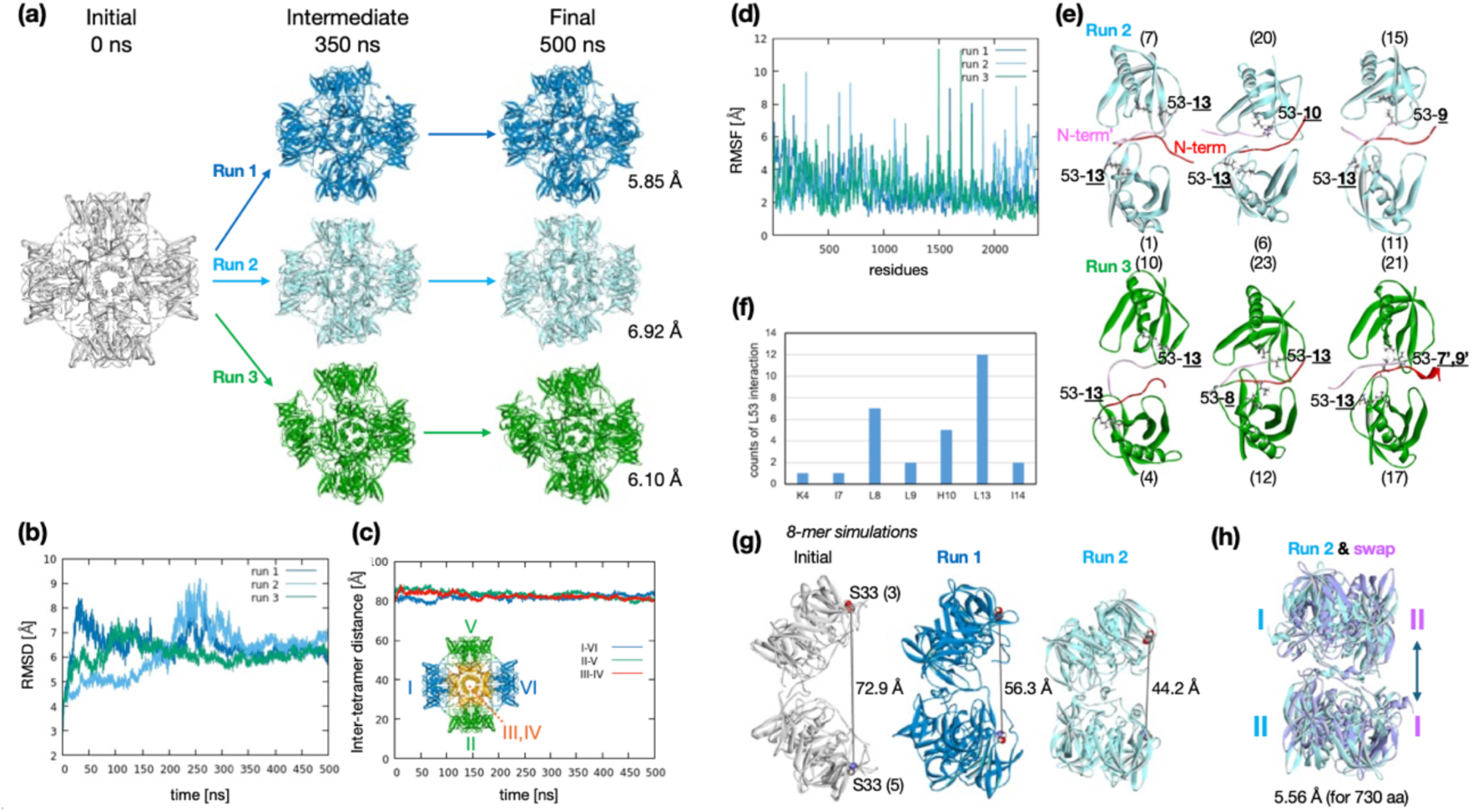
MD simulations of 24-mer and 8-mer CipB. (a) 24-mer CipB structures in MD simulations. Runs 1, 2 and 3 are shown in blue, cyan, and green, respectively. (b) RMSDs of 24-mer CipB for 500 ns. (c) Inter-tetramer distance between I-VI, II-V, III-VI tetramer pairs (i.e., between the four L68 mass centers in the tetramer pair) in run 3. (d) RMSFs of 24-mer CipB for 2,400 residues. (e) Structures of six selected inter-tetramer pairs and Leu53 interactions with the N-terminal region. L53 and interacting residues are shown as sticks, and N-terminal regions are shown as red and pink ribbons. (f) Frequency (counts) of Leu53 interactions with N-terminal residues at 500 ns. (g) 8-mer CipB structures in MD simulations and the S33-S33 distances of the furthest monomers. S33 residues are shown as spheres. (h) Superimposed structures of 8-mer CipB (run 2) and its swapped structure prepared using TM-align.(22)

Next, to examine possible assembly intermediates related to the low-molecular-weight oligomers detected by Native-PAGE analysis and cryo-EM (Figs. S3 and S5f), we performed three independent 500-ns MD simulations of an 8-mer model (a dimer-of-tetramers, ∼90 kDa) to explore whether such a subassembly could maintain tetramer-tetramer contacts. In all three simulations, the dimer-of-tetramers was maintained, although the arrangements were found to differ between runs. In runs 1 and 2, the tetramers adopted a relatively compact conformation, whereas in run 3, the tetramers underwent significant rotational rearrangement while maintaining contacts between the N-terminal regions (Fig. S7b). The RMSDs were found to be approximately 12 Å in runs 1 and 2, and 18 Å in run 3 (Fig. S7c). In the compact conformation observed in run 2, both tetramers were found to rotate counter-clockwise in an approximately symmetric manner (Fig. S7d). The Ser33-Ser33 distance between the most distant monomers was found to decrease from an initial value of 72.9 Å to 56.3 Å in run 1 and 44.2 Å in run 2, indicating the formation of more compact conformations, whereas the distance was found to increase slightly to approximately 80 Å in run 3, consistent with a more open arrangement (Figs. 5g and S7e).

Analysis of the N-terminal conformations in the two compact assemblies (run 1 and 2) revealed that contacts between the N-termini of the tetramers gradually increase during the simulations. Initially, only one pair of N-termini from monomers 1 and 7 were found to interact with each other. In run 1, three N-termini from one tetramer (monomers 1, 2, and 4) formed contacts with two N-termini from the other tetramer (monomers 6 and 7) (Fig. S7f). In run 2, three N-termini from each tetramer (monomers 1, 2, 4 and monomers 6, 7, 8) formed more extensive inter-tetramer interactions, consistent with stabilization of the compact conformation (Fig. S7g). The most compact assembly observed in run 2 was found to exhibit a quasi-symmetric arrangement of the two tetramers upon structural superposition of the tetramer-swapped assembly (Fig. 5h, RMSD: 5.56 Å for 730 aa by TM-align(22)). These simulations suggest that the flexible N-terminal region contributes to local inter-tetramer contacts, allowing the components of the dimer-of-tetramers to remain associated with each other while testing different compact and open arrangements.

### Mutation analysis of N-terminal and inter-cage contacts required for CipB lattice formation

The N-terminal region (Asp6–Val15), which is not visible in the crystal structure, was observed between tetramers within a 24-mer cage structure in solution by cryo-EM. MD simulations further suggest that the N-termini participate in intra-cage and inter-tetramer contacts. These observations indicate that the N-terminal region is locally flexible and structurally heterogeneous, while being positioned to participate in contacts between tetrameric units. We therefore examined whether the N-terminal region is required for CipB lattice formation in cells.

To test this model, we constructed N-terminal truncation mutants (**CipB_Δ2– 12**, **CipB_Δ2–16**, and **CipB_Δ2–19**). All three were expressed but were recovered almost exclusively from the insoluble fraction. Even the shortest truncation, **CipB_Δ2– 12**, failed to produce detectable intracellular particles, whereas the longer truncations formed amorphous aggregates (Fig. S8). None of the mutants released the soluble 24- mer species observed for wild-type CipB upon incubation across pH 3.0-10.0 (Fig. S9), and SAXS measurements showed neither Bragg peaks nor the scattering features characteristic of the hollow cage (Figs. 1d and S10a). These results show that the N-terminal region is required for CipB crystallization in cells, although the accompanying loss of solubility means they do not isolate a specific requirement for cage assembly from a more general destabilization of the fold.

To investigate the effect of inter-cage interactions in the CipB crystal lattice formation, we introduced site-specific mutations at Val46, which mediates hydrophobic contacts between neighboring cages in the crystal structure. Substitution with leucine (**CipB_V46L**) or isoleucine (**CipB_V46I**) yielded in-cell crystals and exhibited SAXS diffraction patterns consistent with a bcc lattice with a 111.3 Å lattice constant, which is similar to that of the wild type (Fig. S10b). In contrast, substitution with serine (**CipB_V46S**) eliminated crystallization, as indicated by the absence of diffraction peaks in SAXS measurements (Fig. S10b). These findings suggest that V46 contributes to crystal lattice assembly by promoting appropriate hydrophobic interactions between neighboring cages.

To further assess the role of residues at the inter-cage interface, we introduced point mutations at His75 and Lys77, which form side-chain contacts between adjacent cages in the crystal structure. Several variants, including **CipB_H75T**, **CipB_H75Y**, **CipB_K77A**, and **CipB_K77Q**, retain the ability to crystallize in cells and exhibit SAXS diffraction profiles consistent with a bcc lattice, matching the parameters of **CipB_icc** (Fig. S10c). These mutational analyses indicate that the N-terminal region and hydrophobic Val46-mediated inter-cage contacts are required for SAXS-detectable CipB lattice formation, whereas His75 and Lys77 are not individually essential. This may reflect the capacity of variant residues at these positions to form alternative non-covalent interactions at this interface, or the contribution of backbone geometry in the His75–Lys77 region to inter-cage packing independently of specific side-chain identity.

## Discussion

In this study, we analyzed the structures of CipB in both crystal and solution states to understand the molecular basis of in-cell crystallization of CipB. Our results show that CipB forms a stable 24-mer cage in solution, consistent with its role as a building block for crystal lattice formation. In particular, the structurally flexible N-terminal region contributes to contacts within the cage assembly. Structural analyses by crystallography, cryo-EM, and MD simulations consistently revealed local flexibility in the N-terminal and surface-loop regions, while the overall cage structure is maintained. These findings suggest that structurally flexible supramolecular building blocks may facilitate in-cell crystallization by accommodating local conformational variability. The key feature of CipB crystallization is not simply the formation of a 24-mer cage, but the coexistence of a preserved cage architecture with local conformational flexibility at the interfaces that mediate higher-order assembly.

Our results support for a stepwise assembly model for CipB crystallization in living cells (Fig. 6a). The observation that **CipB_sol** reforms the same bcc lattice upon recrystallization, together with the loss of intracellular crystallization in the N-terminal deletion mutants, supports a model in which the soluble 24-mer serves as a precursor for in-cell crystallization. In this model, the flexible N-terminal region appears to link neighboring tetramers and appears to support the formation of a 24-mer cage. Following formation of the 24-mer cage, hydrophobic interactions between adjacent cages promote packing into the crystal lattice. Val46 is one of the residues contributing to this interface. The cage architecture may accommodate local structural variability while preserving the organization required for crystal lattice formation.

**Fig. 6.**
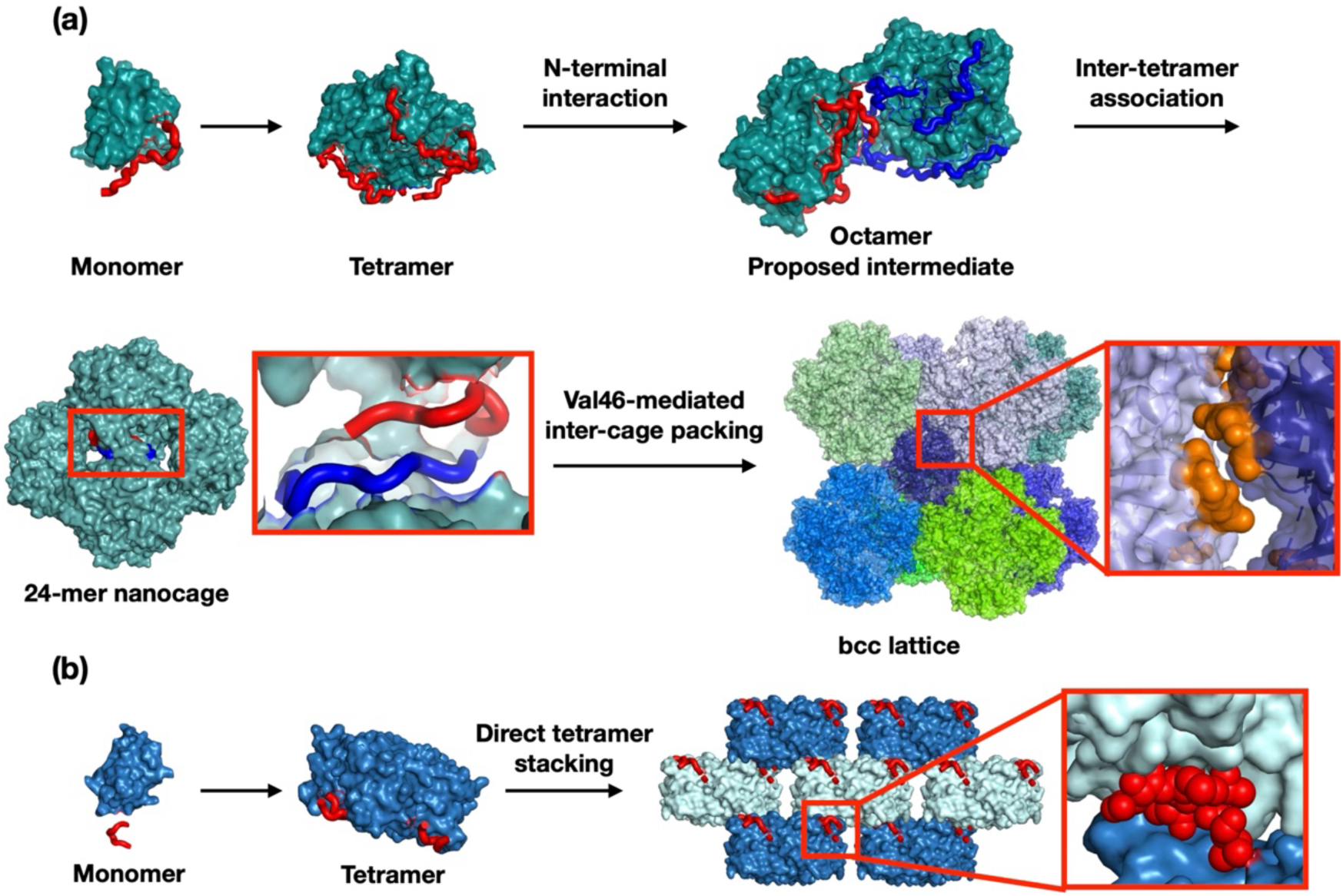
Proposed mechanisms of CipB and CipA crystallization based on structural analyses. (a) CipB assembles from tetramers into a 24-mer assembly through N-terminal interactions via a proposed 8-mer intermediate, followed by bcc lattice formation via Val46-mediated inter-cage packing. (b) CipA crystallizes through direct tetramer stacking.

A notable feature of this assembly pathway is that local structural flexibility is retained even after assembly into the 24-mer cage and crystal lattice. Analysis of normalized *B*-factors derived from the crystal structure suggests that the relatively flexible regions are localized to the cage surface and inter-tetramer interfaces, whereas the β-sheet-rich core regions remain rigid (Fig. S11). Comparison of the crystal and cryo-EM structures further showed that these conformational differences occur in the N-terminal and loop regions located on the cage surface, within the cage interior, and at the inter-tetramer interface (Fig. S6). Cryo-EM analysis further revealed weak or heterogeneous map density in several loop regions of the cage surface and tetramer interfaces. Furthermore, the presence of lower-molecular-weight oligomeric species detected by native PAGE and cryo-EM suggests that the CipB 24-mer cage is not completely homogeneous in solution (Figs. S3 and S5f). These species may represent partially assembled or partially dissociated states associated with the dynamic assembly behavior of the 24-mer cage.

The importance of the flexible N-terminal region is further highlighted by comparison with CipA, which shares approximately 23% amino acid sequence identity and has a similar monomeric structure (Cα RMSD: 1.8 Å).(21) Although both proteins are OB-fold monomers that form tetramers, they exhibit fundamentally different processes for lattice assembly. While CipA directly assembles into a tetragonal lattice through tetramer stacking mediated by N-terminal interactions, CipB stabilizes a 24- mer cage structure in solution via tetramer–tetramer interactions mediated by the flexible N-terminal region (Asp6-Val15). Structural comparisons suggest that differences in the orientation of the N-terminal region are associated with the distinct assembly pathways of CipA and CipB. In CipA, the N-terminal region extends outward from the tetramer, thereby stacking between tetramers within the crystal lattice (Fig. 6b). In contrast, the N-terminal region of CipB bends toward adjacent tetramers, forming pairwise interactions within the 24-mer cage (Fig. 6a). These differences suggest that variations in N-terminal interactions are associated with the distinct lattice formation of CipA and CipB.

This flexible cage contrasts with canonical rigid protein nanocages such as the ferritin cage (PDB 6RJH).(23) While ferritin forms a highly stable 24-mer cage through extensive inter-subunit interactions, CipB retains local flexibility, despite adopting a similar 24-mer architecture. The buried surface area (BSA) per ferritin monomer (3,770 Å^2^, 41% of the solvent-accessible surface area) is significantly greater than that of CipB (1,920 Å^2^, 32%), suggesting weaker inter-subunit interactions within the 24-mer cage of CipB. Cryo-EM and MD simulations further support the retention of conformational flexibility in the N-terminal and surface-loop regions of CipB even after assembly into the 24-mer cage. Unlike ferritin, CipB does not appear to require complete rigidification prior to crystallization. This feature also distinguishes CipB from recently reported computationally designed in-cell protein assemblies, highlighting a natural in-cell crystallization system in which a stable 24-mer assembly retains local conformational flexibility during crystallization.(24)

Furthermore, we successfully produced CipB crystals using a cell-free protein crystallization method (Fig. S12).(21) This demonstrates that CipB crystallization is not limited to the intracellular environment and can also occur under cell-free conditions. These observations suggest that the ability to crystallize is primarily inherent to the protein itself. These findings suggest a design concept in which flexible structural elements can contribute to supramolecular assembly and crystallization processes.

In conclusion, our structural analyses show that CipB crystallizes through a 24-mer assembly precursor instead of undergoing direct packing of monomers or tetramers. The flexible N-terminal regions support a productive association between tetramers within the cage, whereas defined contacts between cage surfaces impose long-range crystalline order. These findings identify three elements that are important for CipB intracellular crystallization: formation of a discrete higher-order precursor, retention of local conformational flexibility within that precursor, and specific recognition between assembly surfaces. Thus, CipB provides a structural example in which conformational flexibility retained within a protein assembly can be maintained during the formation of the ordered lattice of an in-cell protein crystal. By defining these molecular features, this study establishes a basis for understanding and engineering in-cell protein crystallization for future structural and biomaterials applications.

## Materials and Methods

Detailed Materials and Methods are described in *SI Appendix*.

## Data availability

The atomic coordinates of CipB from the crystals have been deposited in the Protein Data Bank under accession code 25YU. The cryo-EM map of the CipB 24-mer cage has been deposited in the Electron Microscopy Data Bank (EMDB) under accession code EMD-81404. The atomic coordinates of the CipB 24- mer cage have been deposited in the Protein Data Bank under accession code 27TD.

## Acknowledgments

Synchrotron radiation experiments were conducted at SPring-8 BL32XU under proposal Nos. 2021A2541, 2021A2744, 2021B2744, 2022A2735, 2022B2735, 2022A2770, and 2022B2770. We thank Dr. Kunio Hirata and the beamline staff at BL32XU of SPring-8 (Sayo, Japan) for their assistance with X-ray crystallographic data collection and processing using the KAMO system. This work was supported by the Japan Society for the Promotion of Science (JSPS) KAKENHI Grant Nos. 22H00347, 25H02254 to T.U.; 22K19266, 25K08825 to S.A; and 26K17957 to K.K. This work was also supported by the Japan Science and Technology Agency (JST) through the Adaptable and Seamless Technology Transfer Program through Target-driven R&D (JPMJTR20U1, JPMJTR224A) to T.U. and ACT-X (JPMJAX25D5) to K.K. This research was partially supported by the Platform Project for Supporting Drug Discovery and Life Science Research [Basis for Supporting Innovative Drug Discovery and Life Science Research (BINDS)] from the AMED under Grant number JP21am0101070 (support No. 1854). ChatGPT (OpenAI) was used for the sole purpose of language editing and improving the clarity of the manuscript.

