## Supporting information for "In-Cell Protein Crystallization via a Locally Flexible 24-mer Assembly Precursor"

### Table of Contents

**Experimental Procedures**

| | |
|---|---|
| **Materials** | 3 |
| **Plasmid Construction and Expression** | 3 |
| **Identification of in-cell protein crystals of CipB (CipB_icc)** | 3 |
| **Small-angle X-ray Scattering (SAXS) Measurement** | 4 |
| **Preparation and identification of CipB_sol** | 4 |
| **Dynamic Light Scattering (DLS) measurement** | 5 |
| **Recrystallization and X-ray crystal structural analysis** | 6 |
| **Cryo-EM grid preparation, data collection, image processing and model building** | 6 |
| **MD simulations** | 8 |
| **Construction and characterization of CipB mutants** | 8 |
| **Cell-free crystallization of CipB** | 9 |

**Supplementary Figures and Tables**

| | |
|---|---|
| **Fig. S1** | 10 |
| **Fig. S2** | 11 |
| **Fig. S3** | 12 |
| **Fig. S4** | 13 |
| **Fig. S5** | 14 |
| **Fig. S6** | 16 |
| **Fig. S7** | 17 |
| **Fig. S8** | 18 |
| **Fig. S9** | 19 |
| **Fig. S10** | 20 |
| **Fig. S11** | 21 |
| **Fig. S12** | 22 |
| **Table S1** | 23 |
| **Table S2** | 24 |
| **Movie S1** | 25 |
| **Supplementary References** | 25 |

#### Experimental Procedures

##### Materials

The *E. coli* cells BL21 (DE3) provided by Invitrogen were used as the standard strain. The plasmid used in this study is pET29 from Merck. Other chemicals were purchased from Wako, Nacalai Tesque, Sigma-Aldrich, Qiagen, and Takara Bio and were used without further purification.

##### Plasmid Construction and Expression

The CipB gene was cloned into the pET29 vector for expression in *E. coli*. The plasmid was amplified in DH5α bacteria and purified using the Qiagen Plasmid Mini Kit, and then transformed into *E. coli* BL21 (DE3) cells, which were grown in LB medium containing 30 μg/ml kanamycin and 0.05% glucose. Cells were pre-cultured at 37 °C until $OD_{660}$ reached 0.6-0.8. The pre-culture was inoculated into fresh LB medium containing 30 μg/mL kanamycin, and expression was induced by adding 1.0 mM isopropyl β-D-1-thiogalactopyranoside (IPTG), followed by a 16- to 20-hour incubation at 30 °C with shaking. Cells were harvested by centrifugation at 1,600 g for 5 minutes at 4 °C. The cells were washed with PBS buffer and then disrupted by sonication. Next, the crystals were collected by centrifugation at 1,150 g for 3 minutes at 4 °C, then washed with PBS. Finally, the crystals were stored at 4 °C in PBS.

##### Identification of in-cell protein crystals of CipB (CipB_icc)

The purified **CipB_icc** were identified by SDS-PAGE, Matrix-assisted laser desorption ionization-time of flight mass spectrometry (MALDI-TOF MS), and SEM. A suspension of crystals in PBS was mixed at a 1:1 volume ratio with SDS sample buffer containing 0.17 M DTT, and then boiled for 5 minutes. The samples were separated in an SDS-PAGE gel and stained with the CBB staining solution provided by Nacalai.

For MALDI-TOF MS analysis, sinapinic acid (purchased from Sigma) was used as the matrix. We applied the double-layer method for sample preparation; the first layer consisted of 2 μL of saturated sinapinic acid in ethanol deposited on the target plate, and the second layer consisted of 2 μL of sample in TA solution (1 μL of suspended crystal mixture + 5 μL of a 30:70 (v/v) mixture of acetonitrile and 0.1% trifluoroacetic acid) deposited onto the first layer. The molecular mass of CipB was determined by the Ultrafle Xtreme system (Bruker Daltonics). SEM measurement using the JEOL JCM-6000 system confirmed the morphology of purified crystals. After substituting PBS with Milli-Q water, the crystals were dried, coated with gold, and observed by SEM. Crystal size distributions were analyzed, and histograms were generated using ImageJ software.

**Small-angle X-ray Scattering (SAXS) Measurement**

SAXS measurements were performed using MicroMax-007HF (Rigaku), a microfocus rotating anode X-ray generator equipped with a Cu-Kα source providing the wavelength λ = 1.54 Å. The scattered X-ray intensity was collected by the detector Pilatus 100K-S (Rigaku). The sample-to-detector distance was set to 706.5 mm. Protein crystals were resuspended in a relatively high concentration and filled into a 1.5 mm-thick cell. The 1D scattering profile I(q), plotted as a function of the scattering vector q, was analyzed using the Smartlab Studio II software (Rigaku).

**Preparation and identification of CipB_sol**

In-cell crystals of CipB (**CipB_icc**) purified from *E. coli* were suspended in 100 mM buffer solutions at various pH values: sodium citrate-citric acid (pH 3.0 and 6.0), sodium acetate-acetic acid (pH 4.0 and 5.0), HEPES-NaOH (pH 7.0 and 8.0), Gly-NaOH (pH 9.0), or sodium carbonate-sodium bicarbonate (pH 10.0). Suspensions were incubated at room temperature for

24 h. The resulting supernatants were analyzed by native PAGE. Each supernatant was mixed at a 1:1 (v/v) ratio with a 0.1% BPB solution and loaded onto a 7.5% polyacrylamide separating gel.

The supernatant obtained from CipB_icc, dissolved in 100 mM sodium acetate buffer (pH 5.0), was further purified using a Sephadex G-200 gel filtration column. The peak was collected and concentrated using Amicon Ultra centrifugal filters (10 kDa cutoff, Merck).

For SAXS analysis, the concentrated purified CipB (**CipB_sol**) (62 μM) was subjected to measurement, and the scattering curve was fitted using the core-shell model in the SasView software package.

For ESI-TOF MS analysis, **CipB_sol** (122 μM) was measured in 100 mM ammonium acetate (pH 5.0) using LCT Premier (Waters). The observed mass of the 24-mer (275,565 Da) deviated by ~1.5% from the calculated value (271,574 Da), which is within the typical accuracy range for native ESI-TOF MS of large protein assemblies and is likely attributable to buffer adducts and charge-state assignment uncertainty at high m/z.

**Dynamic Light Scattering (DLS) measurement**

DLS measurements were performed using a Zetasizer μV system (Malvern). A protein solution in deionized water, PBS (at pH 7.4, prepared from tablets (T9181, Takara)), or 100 mM buffer (pH 3.0 and 6.0: sodium citrate-citric acid, pH 4.0–5.0: sodium acetate-acetic acid, pH 7.0–8.0: HEPES-NaOH, pH 9.0: Gly-NaOH, pH 10.0: sodium carbonate-sodium bicarbonate) was used for the experiment. 25 μL of the sample was introduced into the ZEN2112 cuvette (Malvern) and equilibrated at 25 °C for 10 s. The means were calculated from four measurements using the averaging function of the Zetasizer software and presented as a volume mean diameter ± s.d. of four measurements.

**Recrystallization and X-ray crystal structural analysis**

Crystallization of **CipB_sol** was performed by the hanging-drop vapor diffusion method. Initial crystallization screening was conducted using the Hampton Research Crystal Screen kit, and the optimal reservoir condition was identified as 0.1 M MES buffer (pH 6.5), 0.01-0.3 M CsCl, and 20% Jeffamine M-600. The crystallization drops were prepared by mixing with 1.0 μL of concentrated **CipB_sol** (24-32 mg/mL) and 1.0 μL of reservoir solution. The mixtures were equilibrated against the reservoir solution at 20 °C. Crystals appeared within one week. Before data collection, the crystals were immersed in a reservoir solution containing 25% ethylene glycol, mounted on MicroLoops (Mitegen), and then frozen in liquid nitrogen. The diffraction data of the CipB crystals were collected at 100 K on the beamline of BL32XU at SPring-8 using an X-ray wavelength of 1.00 Å. Multiple small-wedge datasets were obtained with the automated data-collection system ZOO.(1) The datasets were automatically processed and merged with KAMO,(2) which employs the XDS package.(3) To select isomorphic datasets for merging, KAMO performed hierarchical clustering based on pairwise intensity correlation coefficients.(4) The cluster with the highest $CC_{1/2}$ was used for subsequent analyses. The structure was solved by molecular replacement using *Phaser-MR*(5) from the *CCP4* suite with CipA (PDB 7XHS) as the search model.(6) Rebuilding was completed using *COOT(7)* based on sigma-A weighted $2|F_o|-|F_c|$ and $|F_o|-|F_c|$ electron density maps. The models were subjected to quality analysis at various refinement stages using omit maps and *RAMPAGE.*(8) Buried surface areas of the CipB and ferritin (PDB: 6RJH) were estimated using the Proteins, Interfaces, Structures, and Assemblies (PISA).(9)

**Cryo-EM grid preparation, data collection, image processing and model building**

3 μL of the wild type CipB complex was applied onto a Quantifoil R1.2/1.3 200 mesh Cu grid (Quantifoil, Jena, Germany) that had been pretreated by glow discharge. After sample

application, excess solution was blotted off and the grid was immediately plunged into liquid ethane for vitrification using a Vitrobot Mark IV (Thermo Fisher Scientific, USA). Cryo-EM data collection was performed on a CRYO ARM 300 II transmission electron microscope (JEOL, Japan) operated at an accelerating voltage of 300 kV. Images were recorded using a K3 camera (Gatan Inc., USA). The defocus range during data collection was set from 0.8 to 2.2 underfocus, with a total dose of 40 $e^{-}/Å^{2}$ on the specimen. Data collection was carried out using the SerialEM program.(10) Image processing was performed using the cryoSPARC v4.6.2 software package.(11, 12) A total of 3,851 movie micrographs were subjected to patch motion correction. The contrast transfer function (CTF) of each micrograph was estimated using patch CTF estimation. To perform automated particle picking, reference 2D class averages were first obtained from manually picked particles. Subsequently, 1,714,413 particles were picked from the total micrographs using the template picker. After 2D classification, well-converged CipB complexes were selected, comprising 443,675 particles. This set of particles was reconstructed and refined using *ab initio* reconstruction and homogeneous refinement, respectively. The resultant consensus map showed partial destruction, indicating that this data set contained structural heterogeneity. To isolate structurally intact CipB particles, 3D classification was performed. Among the three classes, the major class displayed intact structural features. This data set was further subjected to homogeneous refinement and non-uniform refinement. The obtained final reconstruction reached a resolution of 3.63 Å. Cryo-EM structures were visualized using the ChimeraX software.(13) The structural model was constructed using UCSF ChimeraX(14) and COOT, with a starting model based on the crystal structure. The atomic model of CipB was built and refined in real space using phenix.real_space_refine routine in the PHENIX package.(15) Graphical figures were prepared using UCSF ChimeraX and PyMOL.

**MD simulations**

Based on the crystal structure obtained by X-ray crystallography in this study, the N-termini of the A-B pairs of the cryo-EM structure (in this study) were first fused to each site (24 sites in total), and the missing residues inside were also supplemented from the cryo-EM structure. Next, the remaining missing N-termini were added as extended conformations from the AlphaFold database structure (AF-P96969, https://alphafold.ebi.ac.uk/), forming a 24-mer cage of CipB with each residue 100 aa. The created 24-mer cage was solvated, and counterions were added. Then, with the main chain restraints (10 kcal $mol^{-1}$ $Å^{-2}$), energy minimization was performed for 300 steps, followed by 500 ps of NVT and NPT equilibration (at three different initial velocities). Finally, three independent 500 ns production runs were performed. In addition, the 8-mer system (a dimer of adjacent tetramers) was subjected to three runs using the same procedure as above. The temperature and pressure were regulated by the Berendsen (weak coupling) thermostat and barostat (300 K and 1 bar), respectively. All MD simulations were performed using the Amber 24 package, with force field parameters for ff19SB (protein) and OPC (water molecule).(16)

**Construction and characterization of CipB mutants**

The CipB mutants (**CipB_Δ2–12**, **CipB_Δ2–16**, **CipB_Δ2–19**, **CipB_V46L**, **CipB_V46I**, **CipB_V46S**, **CipB_H75T**, **CipB_H75Y**, **CipB_K77Q**, **CipB_K77A**) were generated by PCR-based site-directed mutagenesis using the KOD Plus mutagenesis Kit (Toyobo). The mutant proteins were expressed in *E. coli* cells BL21 (DE3) under the same conditions as wild-type CipB. Intracellular crystal formation was evaluated by microscopy and SAXS.

**Cell-free crystallization of CipB**

Expression and crystallization of CipB were performed using a WEPRO7240 Expression Kit (CellFreeSciences).(6) The CipB gene was cloned into the PEU-E01-MCS vector (CellFree Sciences) to express CipB. The plasmid was amplified in DH5α bacteria and purified using the Qiagen Plasmid Midi Kit. Transcription was performed using a dialysis system according to the expression kit protocol. After 6 h of incubation at 37 °C, mRNA was used for translation. Translation reactions were carried out using the dialysis method. 60 μL of reaction mixture containing 20 μL of WEPRO7240 wheat germ extract, 20 μL of prepared mRNA, 20 μL of SUB-AMIX SGC solution, and 40 mg/mL creatine kinase was dialyzed against 2.5 mL of SUB-AMIX SGC solution in a cylindrical cup and incubated at 20 °C for 72 h. The crystals were collected by centrifugation and washed several times with PBS.

**Supplementary Figures and Tables**

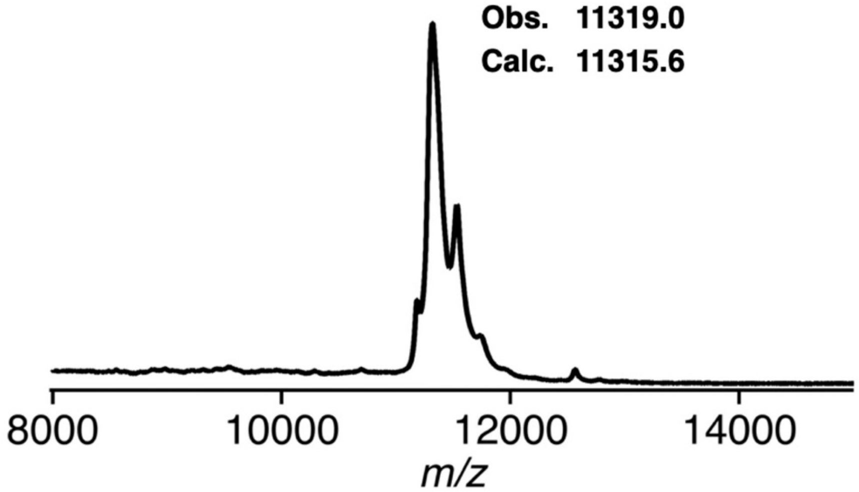


**Fig. S1.** MALDI-TOF mass spectrum of purified **CipB_icc**.

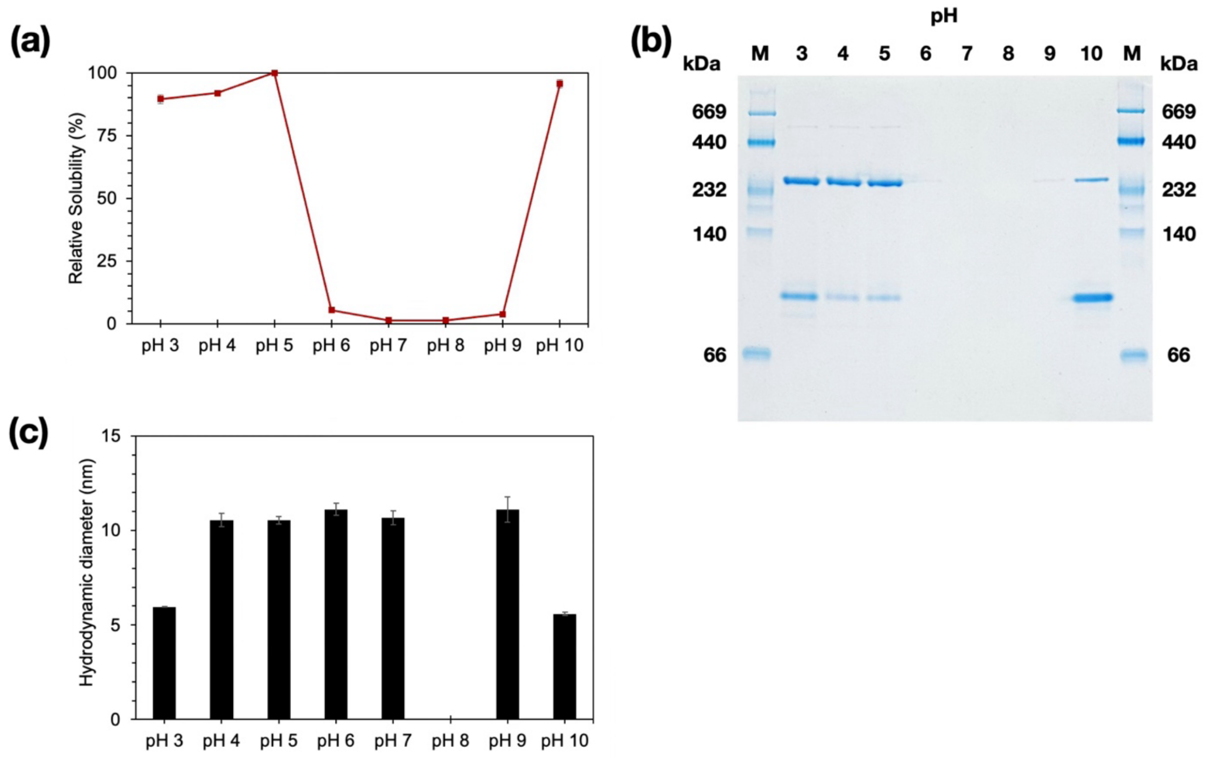


**Fig. S2**. (a) Relative solubility of **CipB_icc** after incubation in buffers ranging from pH 3.0 to 10.0, (b) Native PAGE analysis of the supernatant fractions after incubation of **CipB_icc** in the same buffer (M: marker). (c) Dynamic light scattering (DLS) measurements of the hydrodynamic diameter of the corresponding supernatant fractions. **CipB_icc** was incubated at room temperature for 24 h in 100 mM buffers of the following compositions: sodium citrate-citric acid (pH 3.0 and 6.0), sodium acetate-acetic acid (pH 4.0 and 5.0), HEPES-NaOH (pH 7.0 and 8.0), Gly-NaOH (pH 9.0), or sodium carbonate-sodium bicarbonate (pH 10.0). No DLS signal was detected at pH 8.0 because CipB_icc did not dissolve.

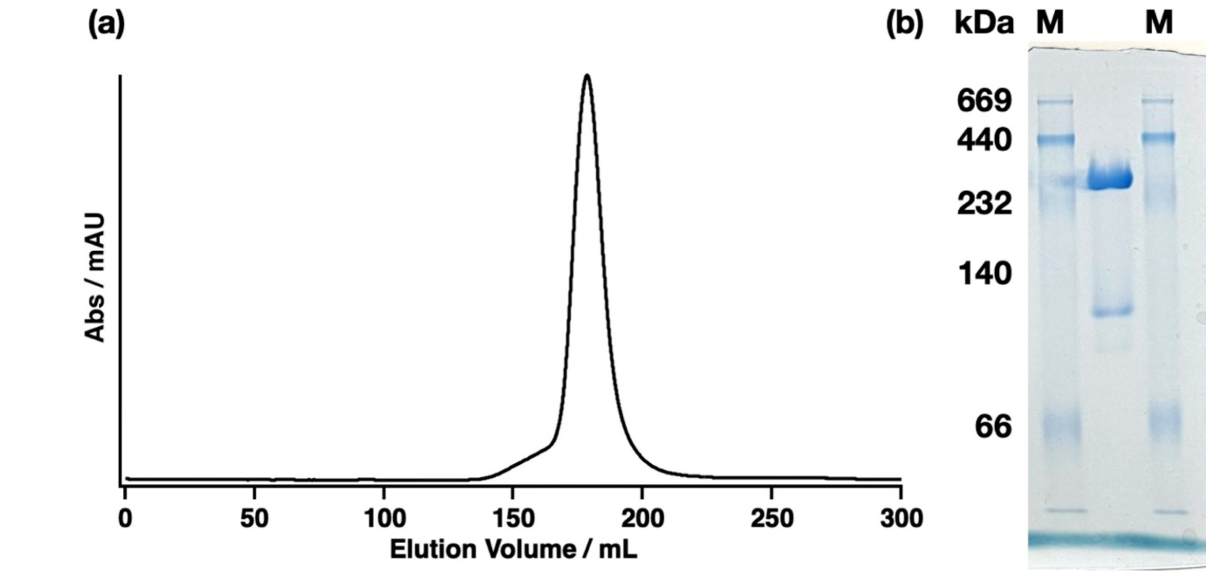


**Fig. S3.** (a) Size exclusion chromatography (SEC) profile of dissolved **CipB_icc** at 100 mM sodium acetate buffer (pH 5.0). The peak was collected and used for SAXS measurements, as shown in Figure 1d. (b) Native PAGE of the collected fractions confirmed the presence of consistent oligomeric species across the peak, indicating that the assemblies observed by SAXS represent the dominant solution species.

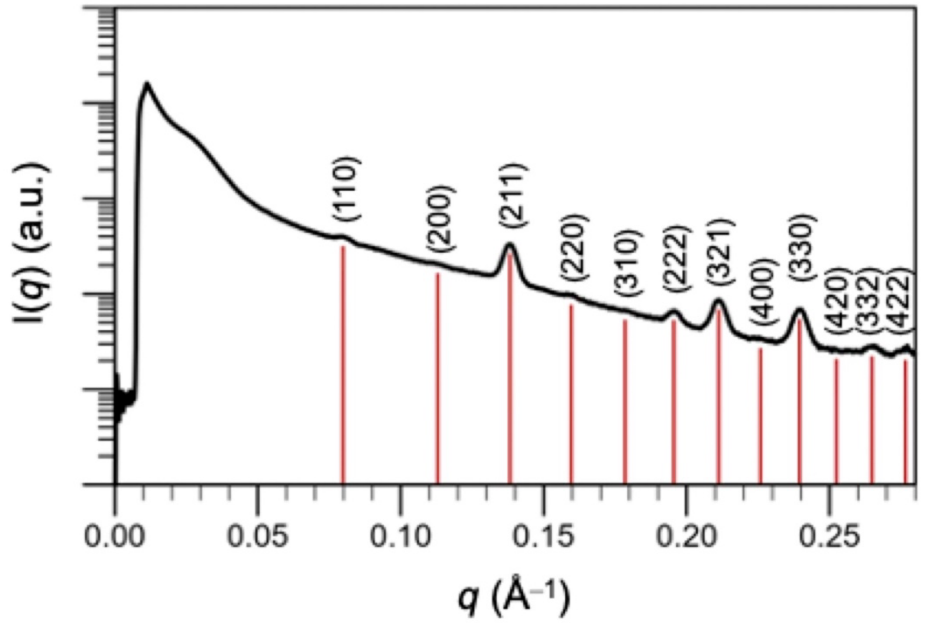


**Fig. S4.** SAXS profile of the purified **CipB_icc**, indicating a body-centered cubic (bcc) lattice with characteristic diffraction peaks. The simulated pattern for the bcc lattice (a = 111.3 Å) is shown as a red line.

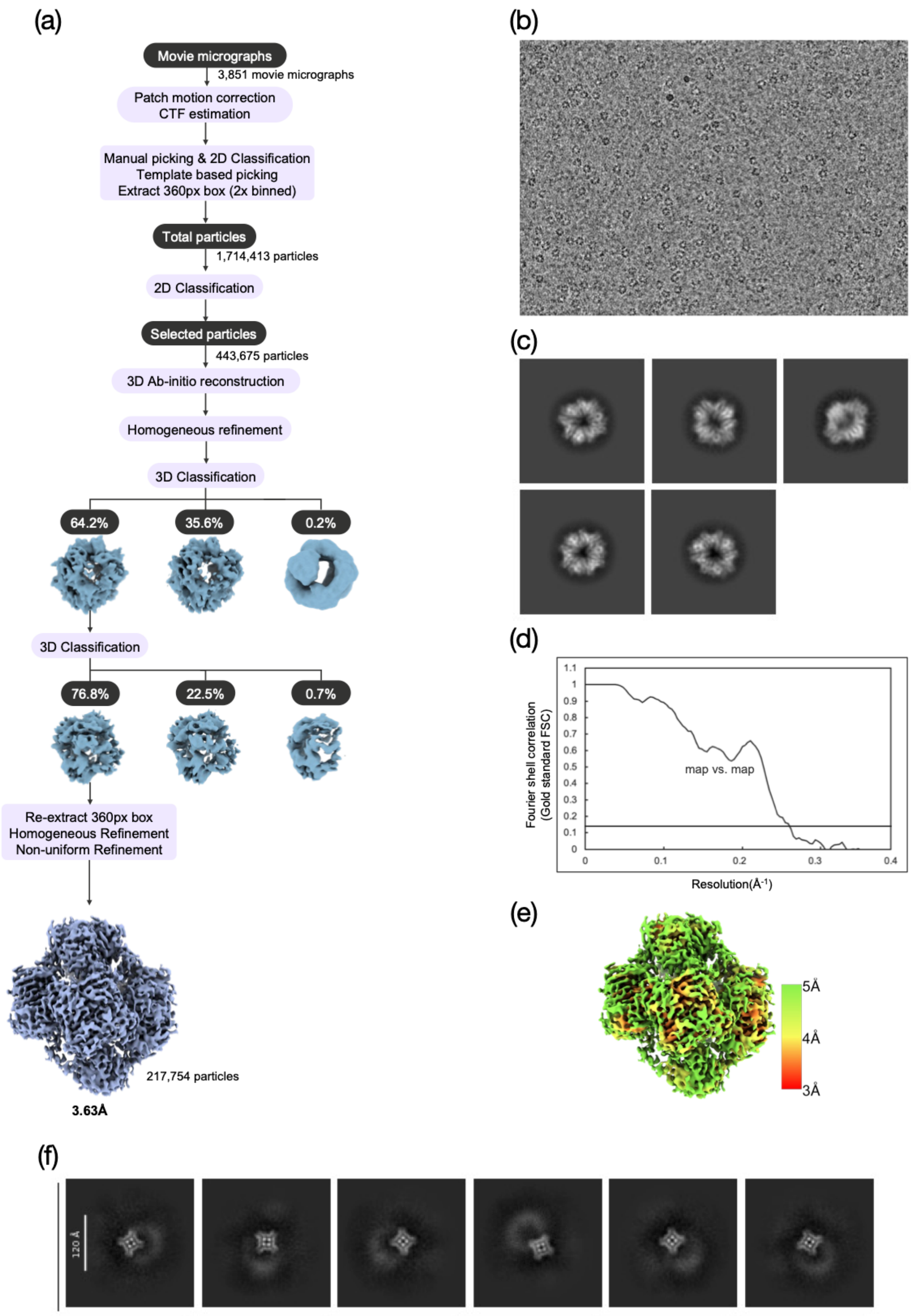


**Fig. S5.** Cryo-EM image processing workflow for the CipB_sol. (a) The workflow of single-particle analysis of CipB. A total of 3,851 movie micrographs were motion-corrected. Subsequently, particles of the oligomerized CipB complex were manually picked and 2D classified to generate references for template picking. 1,714,413 particles were extracted and subjected to 2D classification, resulting in the selection of 443,675 CipB complex particles. Ab-initio reconstruction and homogeneous refinement were then performed to obtain a three-dimensional structure. To further improve the reconstruction, 3D classification was carried out. One of the three classified structures retained the intact overall feature, whereas the remaining classes were partially degraded. The major class, comprising 217,754 particles, was further refined, yielding a final reconstruction at an average resolution of 3.63 Å. (b) A representative micrograph and (c) 2D class average of CipB. (d) Resolution curve. The gold standard FSC for the final reconstruction is shown. (e) Local resolution distribution of the map. (f) Representative 2D class averages of small particles that emerged during 2D classification of the CipB 24-mer particles.

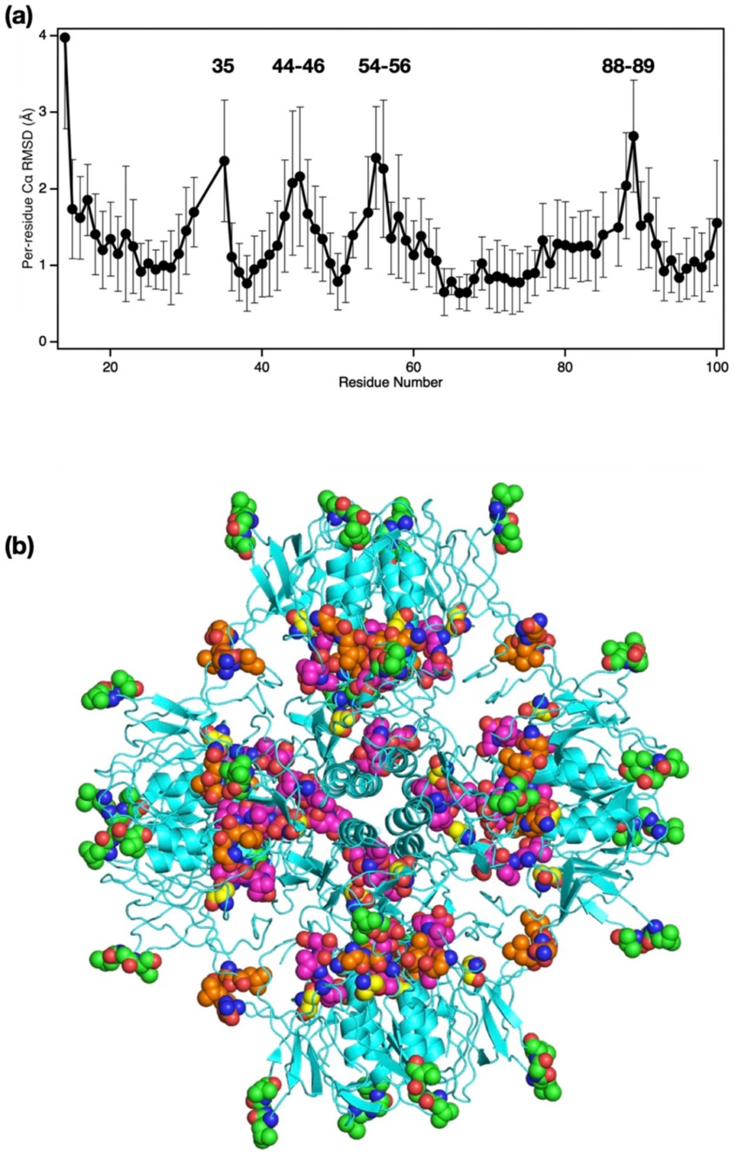


**Fig. S6.** Structural differences between the crystal and cryo-EM structures of the CipB 24-mer assembly. (a) Per-residue Cα RMSDs between the crystal and cryo-EM structures of the CipB monomer. Error bars indicate the standard deviations of Cα deviations among the 24 cryo-EM monomers after superposition to the crystal structure. Regions showing relatively large structural deviations are indicated. (b) Residues exhibiting elevated Cα RMSDs are mapped onto the cryo-EM structure of the CipB 24-mer assembly and are shown as spheres. Ser35 (yellow) and Leu54-Asp56 (magenta) are located within the cage interior, Leu44-Val46 (green) is located on the cage surface, and Asn88-Leu89 (orange) is located at the inter-tetramer interface.

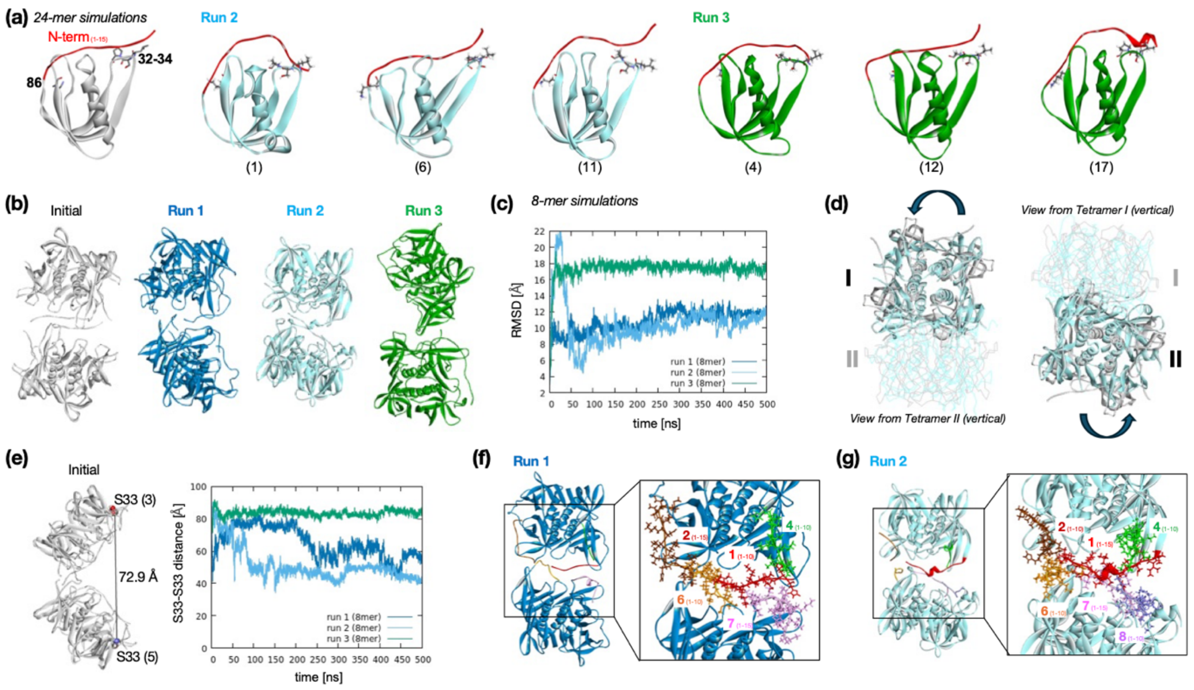


**Fig. S7**. N-terminal interactions in 24-mer simulations and MD simulations of 8-mer CipB. (a) Selected monomer structures in 24-mer simulations. Residues Leu32-Pro34 and Lys86 are shown in sticks, and N-terminal regions are in red ribbons. (b) 8-mer CipB structures in MD simulations. (c) RMSDs of 8-mer CipB for 500 ns. (d) Rotations of tetramers (I and II) in run 2. (e) Ser33-Ser33 distances of the furthest monomers for 500 ns. (f) N-terminal interactions in run 1. (g) N-terminal interactions in run 2.

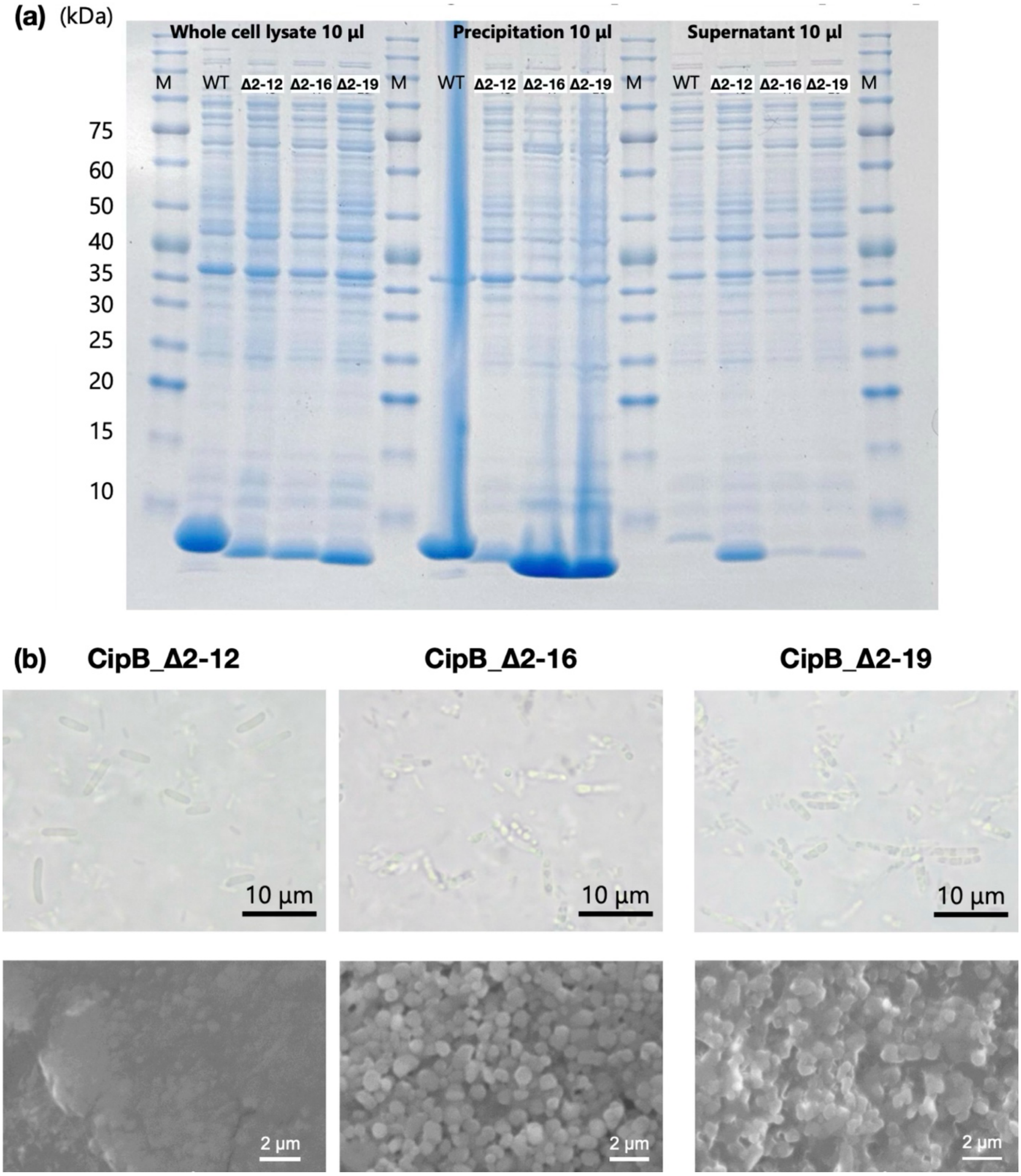


**Fig. S8.** Expression and crystallization behavior of CipB N-terminal truncation mutants. (a) SDS-PAGE confirmed expression of WT and mutants **CipB_Δ 2-12**, **CipB_Δ 2-16**, and **CipB_Δ 2-19** in whole-cell lysates, pellet, and supernatant fractions. (b) Optical microscopy and SEM images show that **CipB_Δ 2-12** failed to form intracellular particles, whereas **CipB_Δ 2-16** and **CipB_Δ 2-19** produced amorphous aggregates instead of ordered crystals.

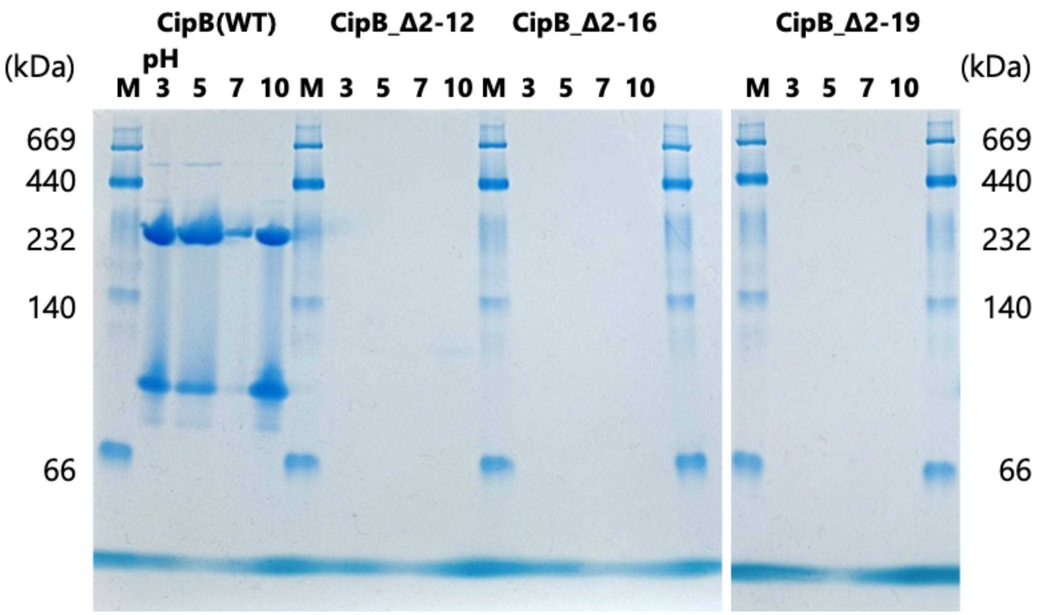


**Fig. S9.** pH-dependent solubilization of CipB WT and N-terminal truncation mutants. Native PAGE analysis of CipB WT and truncated mutants (**CipB_Δ2–12**, **CipB_Δ2-16**, and **CipB_Δ2-19**) following incubation at different pH conditions. WT sample showed soluble bands corresponding to the 24-mer cage, whereas the truncation mutants exhibited little to no detectable soluble species across all conditions, indicating a lack of defined soluble assemblies.

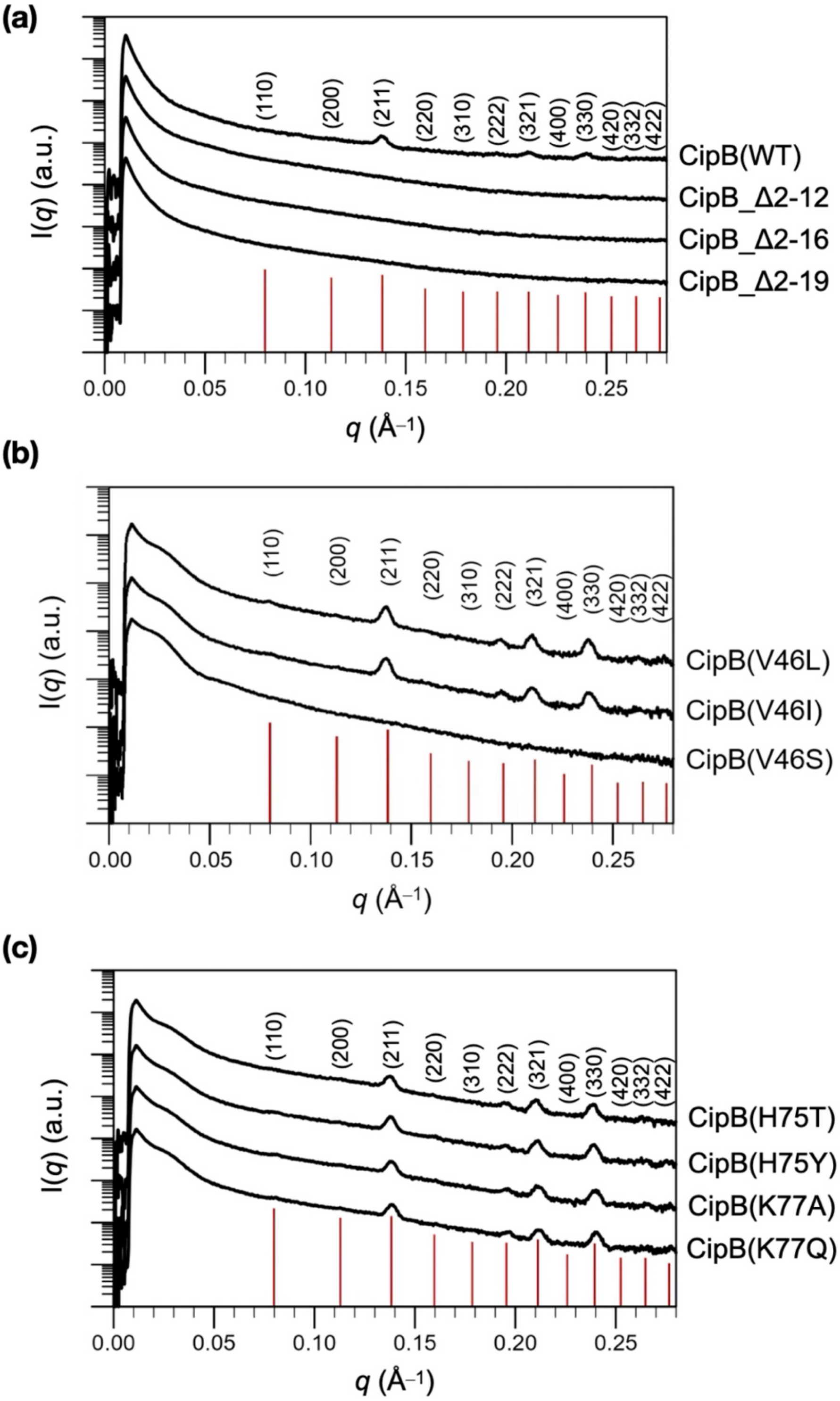


**Fig. S10.** SAXS profiles of CipB variants and truncation mutants. SAXS profiles of (a) N-terminal truncation mutants (**CipB_Δ2-12**, **CipB_Δ2-16**, and **CipB_Δ2-19**), (b) **CipB_V46L**, **CipB_V46I**, and **CipB_V46S** mutants, (c) **CipB_H75T**, **CipB_H75Y**, **CipB_K77Q**, **CipB_K77A** mutants. The samples were prepared as a highly concentrated suspension of *E. coli* expressing CipB mutants in PBS for SAXS measurements.

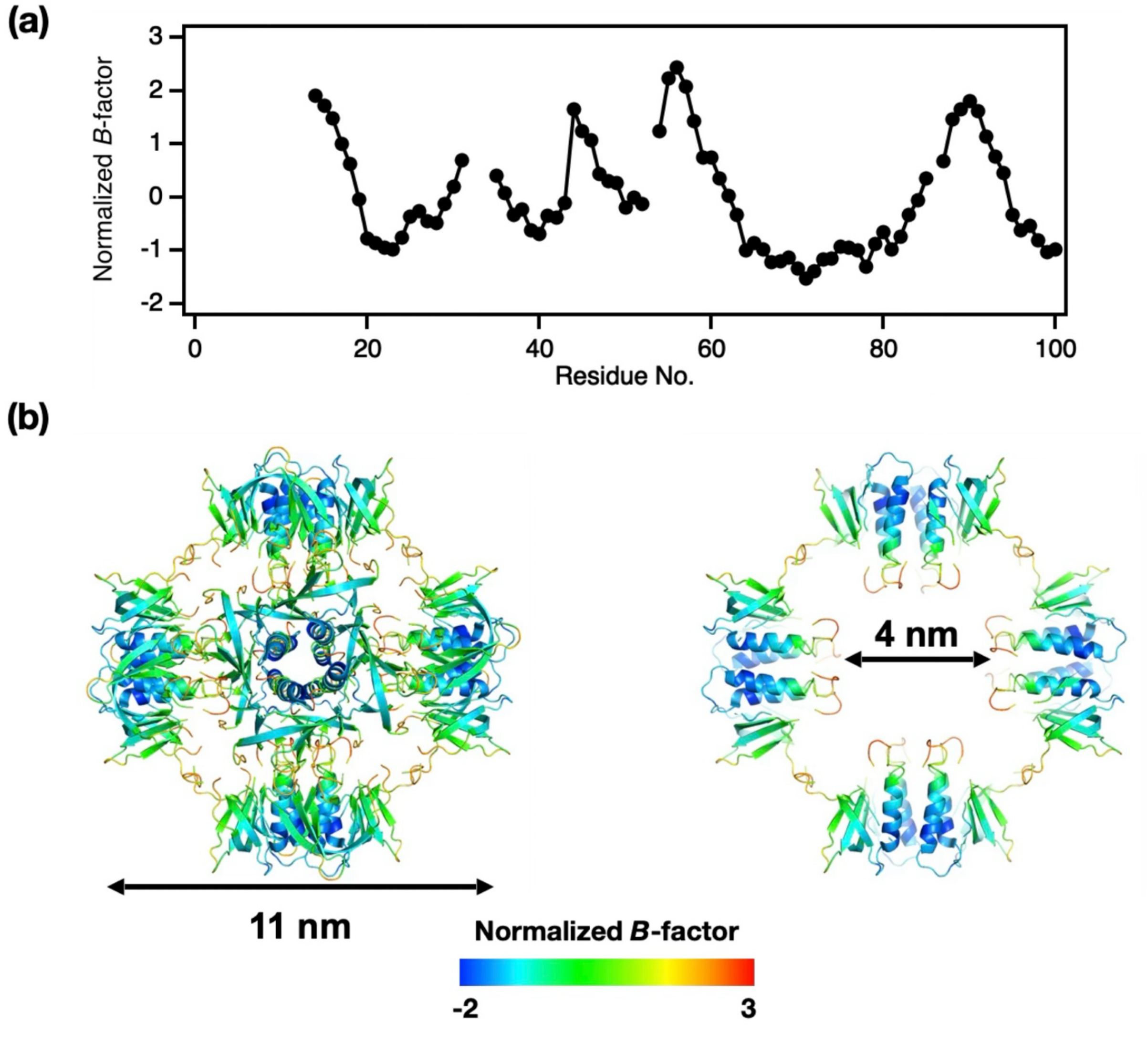


**Fig. S11.** (a) Normalized *B*-factor profile for Cα atoms of CipB. (b) The 24-mer cage structure colored by normalized *B*-factor values (-2 to 3), highlighting flexible loop regions at the cage surface and inter-tetramer interfaces.

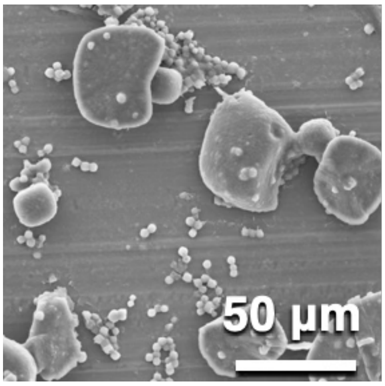


**Fig. S12.** SEM image of CipB crystals with the cell-free protein crystallization method.(6)

**Table S1.** Data collection and refinement statistics of CipB.

| | |
|---|---|
| **Crystallographic data** | |
| Space group | *I*432 |
| Unit cell parameters (Å) | |
| $a = b = c$ | 110.87 |
| Resolution range (Å) | 39.20–3.50 (3.63–3.50) |
| Observations | 86,705 (8,322) |
| Unique reflections | 1,638 (153) |
| Completeness (%) | 100 (100) |
| Multiplicity | 52.9 (54.4) |
| I/σ(I) | 29.3 (8.7) |
| $CC_{1/2}$ | 0.999 (0.983) |
| Rpim | 0.020 (0.130) |
| **Refinement statistics** | |
| Resolution range (Å) | 39.20–3.50 |
| Reflections (work / free) | 1,554 / 82 |
| $R_{work}$ (%) | 32.0 |
| $R_{free}$ (%) | 32.6 |
| R.m.s. deviations from ideal values | |
| Bond lengths (Å) | 0.015 |
| Bond angles (°) | 1.897 |
| Ramachandran plot (%) | |
| Favored region | 83.56 |
| Allowed region | 13.70 |
| Outlier region | 2.74 |

Values in parentheses are for the highest-resolution shell.

**Table S2.** Cryo-EM data collection, model building, refinement, and validation statistics

| | |
|---|---|
| **Data collection and processing** | |
| Magnification | 60k |
| Voltage (kV) | 300 |
| Electron exposure ($e^{-}$/Å$^2$) | 40 |
| Defocus range (μm) | -0.8 to -2.2 |
| Pixel size (Å) | 0.788 |
| Symmetry | C1 |
| Initial particle images (no.) | 443,675 |
| Final particle images (no.) | 217,754 |
| Map resolution (Å) | 3.63 |
| FSC threshold | 0.143 |
| Map sharpening *B* factor (Å$^2$) | -230 |
| **Refinement** | |
| Model composition | |
| Non-hydrogen atoms | 17,562 |
| Protein residues | 2,220 |
| R.m.s deviations from ideal | |
| Bond length (Å) | 0.011 |
| Angle (°) | 1.502 |
| Validation | |
| MolProbity score | 2.97 |
| Clashscore | 40 |
| Poor rotamers (%) | 0.0 |
| Ramachandran plot (%) | |
| Favored region | 64.0 |
| Allowed region | 35.0 |
| Outlier region | 1.0 |

**Movie S1**. Molecular dynamics simulation of the 24-mer CipB (run1).
